# MLO-mediated Ca^2+^ influx regulates root hair tip growth in Arabidopsis

**DOI:** 10.1101/2025.04.08.647801

**Authors:** Sienna T. Ogawa, Weiwei Zhang, Christopher J. Staiger, Sharon A. Kessler

## Abstract

Root hair tip-growth involves coordinated Ca^2+^ and ROS signaling to promote growth while maintaining tip integrity. MILDEW RESISTANCE LOCUS-O (MLO) proteins act downstream of FERONIA (FER) receptor-like kinases in pollen tubes and synergids to regulate calcium dynamics. This study uses a constitutively active MLO (faNTA) to identify a new role for the FER/MLO signaling module in regulating [Ca^2+^]_cyt_ oscillations in growing root hairs.
faNTA bypasses FER signaling in root hairs, complementing not only the previously reported *fer-4* root hair bursting phenotype, but also the complete loss of polarity in spherical root hairs. This complementation correlates with the restoration of tip-focused [Ca^2+^]_cyt_ oscillations that are disrupted in *fer* mutants, suggesting that MLOs are involved in regulating [Ca^2+^]_cyt_ dynamics in growing root hairs.
MLO15 was identified as a regulator of root hair tip-growth based on disrupted root hair growth and [Ca^2+^]_cyt_ signatures in *mlo15-4*. We also link the FER/MLO module to ROS accumulation by showing that faNTA is sufficient to restore ROS levels in *fer-4* root hairs and that ROS levels are decreased in *mlo15-4* root hairs.
We propose that MLOs act downstream of FER to mediate Ca^2+^ influx and promote ROS production in order to regulate root hair tip-growth.

**Plain Language Summary:** Root hairs are important for expanding the surface area of the root for optimal absorption of water and minerals. A FERONIA/MLO signal module promotes tip growth in these specialized cells by regulating calcium and reactive oxygen species accumulation. Understanding tip growth mechanisms may help optimize root responses to environmental challenges.

## Introduction

Plant cells respond to and explore their environments by regulating growth. For example, root hairs are initiated from bulges in epidermal cells and extend out into the soil to maximize water and nutrient absorption. In these tip-growing cells, the site of cell expansion is confined to the apical tip resulting in long cells with a uniform width. Root hair tip growth is modulated through coordination of a number of factors including polarized secretion, cytoskeletal networks, and cell wall modifications (Cole & Fowler, 2006; Mendrinna & Persson, 2015; Takatsuka & Ito, 2020). Furthermore, tip-focused gradients of protons (H^+^), reactive oxygen species (ROS), and cytoplasmic calcium ([Ca^2+^]_cyt_) are associated with polarized growth (Monshausen *et al*., 2008; Stéger & Palmgren, 2022; Lopez *et al*., 2024). In Arabidopsis root hairs, H^+^ oscillations mediated by AHA2 and AHA7 occur at the root hair apex (Monshausen *et al*., 2007; Hoffmann *et al*., 2019). The tip-focused ROS gradient in root hairs is generated by the NADPH oxidase encoded by RESPIRATORY BURST HOMOLOG C/ROOT HAIR DEFECTIVE 2 (RBOHC/RHD2)

(Foreman *et al*., 2003; Monshausen *et al*., 2007). Root hairs also display a tip-focused [Ca^2+^]_cyt_ gradient. [Ca^2+^]_cyt_ oscillations are correlated with root hair elongation (Bibikova *et al*., 1997; Monshausen *et al*., 2008). In root hairs, crosstalk between pH, ROS, and Ca^2+^ has been observed, but how this crosstalk is regulated remains unknown (Monshausen *et al*., 2009).

Hyperpolarized-activated calcium channels (HACCs) expressed at the root tip are thought to mediate the tip-focused [Ca^2+^]_cyt_ gradient and oscillations during tip growth. The presence of HACCs at the root tip was supported by electrophysiology studies, but until recently the identity of these channels remained unknown (Véry & Davies, 2000). Four members of the CYCLIC NUCLEOTIDE GATED CHANNEL family (CNGC5, 6, 9, 14) have been shown to mediate Ca^2+^ influx in root hairs. *cngc14* seedlings have short and branched root hairs when roots are grown into media but appear normal when grown on the surface of vertical plates, indicating that these root hairs are compromised in their ability to respond to the external environment (Zhang *et al*., 2017). When multiple root hair-expressed CNGC genes are knocked out, root hairs are shorter, branched, and frequently burst, indicating problems with maintaining polarized tip-growth (Brost *et al*., 2019; Tan *et al*., 2020). These mutant root hairs have altered [Ca^2+^]_cyt_ oscillations that occur at a lower amplitude and frequency than in the wild type root hairs (Brost *et al*., 2019). Receptor-like kinases are good candidates for regulating Ca^2+^ channels in response to external signals. The *Catharanthus roseus* receptor-like kinase (CrRLK1L) [Ca^2+^]_cyt_-ASSOCIATED PROTEIN KINASE 1 (CAP1)/ERULUS (ERU) functions as a regulator of Ca^2+^ influx in root hairs (Kwon *et al*., 2018). *eru* root hairs are short and have altered [Ca^2+^]_cyt_ oscillations that occur less frequently, but at higher amplitudes that the wild-type (Kwon *et al*., 2018). The Ca^2+^ channels regulated by ERULUS are unknown and it remains unclear whether other families of Ca^2+^ channels contribute to tip growth in root hairs.

The MLO protein family was originally identified as a powdery mildew susceptibility factor in barley (Büschges *et al*., 1997). Since then, members of the MLO family have been implicated in roles throughout the plant including root tropism, pollen tube growth, and pollen tube reception (Chen *et al*., 2009; Kessler *et al*., 2010; Bidzinski *et al*., 2014; Meng *et al*., 2020; Gao *et al*., 2022; Gao *et al*., 2023). Recently, MLO genes in Arabidopsis were characterized as Ca^2+^ influx channels (Gao *et al*., 2022). NORTIA (NTA, MLO7) regulates the amplitude of [Ca^2+^]_cyt_ oscillations in synergids during pollen tube reception (Ngo *et al*., 2014). The CrRLK1L protein FERONIA (FER) is necessary for the initiation of [Ca^2+^]_cyt_ oscillations upon pollen tube arrival at the female gametophyte (Ngo *et al*., 2014). NTA acts downstream of FER: NTA is Golgi-retained before pollination and accumulates in a membrane-rich region known as the filiform apparatus in a FER-dependent manner as the pollen tube approaches (Ju *et al*., 2021). In contrast to *fer* mutants, where synergid [Ca^2+^]_cyt_ oscillations do not occur, *nta-1* synergids have [Ca^2+^]_cyt_ oscillations that occur at reduced amplitude, suggesting that FER likely regulates additional Ca^2+^ channels in synergids (Ngo *et al*., 2014). Our lab discovered that a chimeric protein generated by swapping the C-terminus of NTA with the MLO1 C-terminus (faNTA) accumulates constitutively at the filiform apparatus both before and after pollination (Ju *et al*., 2021). faNTA can bypass FER signaling in synergids and is sufficient to suppress infertility in *fer-1* (Ju *et al*., 2021). These data suggest that faNTA functions as a constitutively activated MLO because it can mediate Ca^2+^ influx at the filiform apparatus independent of FER signaling. In support of this, faNTA accumulates at the plasma membrane in mammalian HEK293 cells and is able to mediate Ca^2+^ influx, while NTA is not plasma membrane localized and instead accumulates in cytoplasmic puncta and is unable to mediate Ca^2+^ influx (Gao *et al*., 2022).

FER is expressed widely throughout the plant and has been implicated in a variety of cellular processes (Ogawa & Kessler, 2023; Cheung, 2024). *fer* mutants have pleiotropic phenotypes ranging from infertility due to a failure of pollen tubes to burst during pollen tube reception to root hairs that burst instead of maintaining tip growth (Escobar-Restrepo *et al*., 2007; Duan *et al*., 2010). The root hair bursting that occurs in *fer* has been attributed to reduced ROS levels due to a failure in FER-mediated ROP-GEF regulation of RBOHC (Duan *et al*., 2010). Since the regulation of root hair tip growth and maintenance of tip integrity has also been linked to the maintenance of [Ca^2+^]_cyt_ oscillations, we hypothesized that FER might also regulate [Ca^2+^]_cyt_ oscillations in growing root hairs through regulation of Ca^2+^ channels. In this study, we test this hypothesis by expressing the constitutively active MLO channel, faNTA, broadly in *fer* mutants and show that it restores tip integrity to *fer* root hairs. We also show that *fer-4* mutants have abnormal [Ca^2+^]_cyt_ oscillations that occur with erratic amplitudes and altered frequency that can be restored to wild-type patterns by expressing faNTA. Furthermore, faNTA is sufficient to restore tip integrity and normal root hair elongation to *fer* mutants, while also restoring ROS to wild-type levels. Finally, we show that MLO15 is expressed in growing root hairs and contributes to root hair elongation by regulating [Ca^2+^]_cyt_ oscillations at the root hair tip. Our study defines a new role for MLO proteins as regulators of [Ca^2+^]_cyt_ oscillations during root hair elongation.

## Materials and Methods

### Plant materials and growth conditions

All seeds were surface-sterilized before plating. *Arabidopsis thaliana* ecotypes Col-0 and Ler were used as wild-type controls. Seeds from these backgrounds were plated on ½ Murashige and Skoog media with 1% sucrose with or without selection. For transgenic selection, the plates were supplemented with hygromycin to a final concentration of 20mg/L. Seeds were stratified at 4 °C for 2 days before being transferred to a growth chamber (22 °C, 16/8 light/dark cycle). The *fer-1* mutant was generously provided by Prof. Ueli Grossniklaus (Escobar-Restrepo *et al*., 2007). Insertion lines for *mlo15-4* (CS66561) and *fer-4* (CS69044) were obtained from the ABRC along with pUBQ10::R-GECO1 in Col-0 (CS72772).

### Cloning and generation of transgenic lines

PCR amplification with Q5 High-Fidelity DNA Polymerase was used to generate the following constructs with Gateway cloning. Entry vector pDONR207-faNTA was described previously (Ju *et al*., 2021). pDONR207-faNTA was recombined via LR reaction into p184 (Myers *et al*., 2016) and pMDC83 (Curtis & Grossniklaus, 2003) with pFER promoter to generate 35S::faNTA-YFP and pFER::faNTA-GFP, respectively. MLO15 was amplified from gDNA with primers MLO15F (GGGGACAAGTTTGTACAAAAAAGCAGGCTTCACCATGGCGGGAGGAG) and MLO15R (GGGGACCACTTTGTACAAGAAAGCTGGGTGATCATGGTGAGCAATCTCT). The MLO15 entry vector was generated via BP reaction with pDONR207 as the backbone. pDONR207-MLO15 was recombined via LR reaction into pMDC83 to generate 35S::MLO15-GFP. pMLO15::MLO15-GFP was previously described (Davis *et al*., 2017). For complementation assays, expression vectors were transformed into *Agrobacterium tumefaciens* GV3101 and transformed into *mlo15-4* and *fer-1/+* using the floral-dip method (Clough & Bent, 1998). Transgenic lines were screened on hygromycin and homozygous T3 or T4 lines were used for experiments.

### Root hair phenotyping, quantification, and live imaging

5 day old vertically grown seedlings were mounted in water and imaged with a Nikon Ti2 microscope. A region of the primary root 2-3 mm above the root tip was chosen for phenotyping because this is the area where *fer* root hair phenotypes are visible and root hairs have not yet burst completely. Root hairs were categorized as tip-growing if they were intact and had polarized growth, burst, or spherical if there was no polarity. Tip-growing root hairs were used for tip width/base width analysis. The tip width was measured as the width of the root hair 10 µm from the tip and the base width was measured at the root hair base. The tip width/base width ratio was determined for all tip growing root hairs and lines were compared with a one-way ANOVA with Tukey test in GraphPad Prism 10.

To quantify root hair length along the entirety of the primary root, seven day old vertically grown seedlings were mounted in water and imaged with a Nikon Ti2 microscope. Root hair length and the distance of that root hair from the root tip was measured using ImageJ (Schindelin *et al*., 2012).

For timelapse imaging of root hairs, seeds were plated on ½ MS with 1% sucrose and stratified for two days at 4 °C. Plates were then grown vertically in a growth chamber (22 °C, 16/8 light/dark cycle) for 5 days. Seedlings were transferred to slides and mounted in liquid 1/10 MS media without sucrose. Parafilm was used to secure the cover slip to the slide and slides were placed vertically in a box with 1/10 MS. The slides were returned to the growth chamber overnight so the seedlings could acclimate before imaging. Root hairs that were closest to the root tip were used for imaging. Images were taken every five minutes for two hours on a Nikon Ti-2 microscope and the imaging room was maintained at 21-22 °C. Root hair length was measured at each time point in ImageJ and the increase in length was plotted over time in Graphpad Prism 10 to obtain average growth rate as the slope of the linear regression.

### Light sheet fluorescence microscopy sample preparation

Seeds were surface sterilized and stratified on ½ MS 1% sucrose plates for at least two days. 100 µL glass capillary tubes (Sigma-Aldrich, BR708744) were cut to the length of the sample holder. Ethylene propylene (FEP) tubing (Bruker Nano, Inc.) with an inner diameter of 1.7 mm was pre-cleaned by washing with 1M NaOH and 70% EtOH as previously described (Weber *et al*., 2014). FEP tubing was cut to 50 mm length and inserted onto the glass micropipette tubes and autoclaved. To prepare samples, the capillary tube/FEP tube was filled with ½ MS, 1% sucrose, 0.5% phytagel medium and a stratified seed was placed on the media at the top of the FEP tube. To keep samples humid and upright, 1mL pipette tips were cut and inserted into sterile jars with 1% agar with a thin layer of water over the top. The capillary/FEP tube with the seed was then inserted into the 1mL pipette tip vertically and the jar covered with a petri dish secured with micropore tape. This system was moved into a growth chamber (22 °C, 16/8 light/dark cycle) and seeds were allowed to grow for five days until root tips were at the bottom of the FEP tubing, but not yet growing through the glass capillary tube.

### Subcellular localization imaging using light sheet and laser scanning confocal microscopy

The complemented line *pMLO15::MLO15-GFP;mlo15-4* was imaged on a Bruker MuVi SPIM light sheet microscope with two Nikon CFI Plan Fluor 10x W 0.3 NA water immersion objective lenses for illumination and two Olympus XLUMPLFLN 20x (eff. 22.2x) W 1.0 NA water immersion objective lenses for detection. The capillary tube/FEP tube with a 5-d-old growing seedling was inserted vertically into the imaging chamber filled with water, keeping the cotyledons out of the water for the duration of imaging. A 2.5 µm beam expander was used. Images were acquired with 10% 488 nm laser intensity, BP 500-530 filter, 50 ms exposure, and a 3 µm step size (175 steps total) every 5 minutes for 4 hours. Images were collected with single side illumination with a Hamamatsu ORCA-Fusion BT CMOS camera. Maximum intensity projections were generated for each time frame in ImageJ.

For laser scanning confocal microscopy, seedlings were incubated in 20 μM FM4-64 (Molecular Probes, T3166) in PIPES Buffer (50 mM PIPES, 5 mM EGTA and 1 mM MgSO_4_ at pH = 6.8) for 5 minutes prior to imaging. Imaging was performed on a Zeiss LSM 880 upright confocal microscope with a 10x objective and 10% laser intensity. Seedings were imaged with a 488 nm excitation wavelength and emission was captured between 485-560 nm for GFP/YFP and 600-850 nm for FM4-64. Image analysis was done in ImageJ.

### R-GECO1 live imaging

R-GECO1 was crossed from Col-0 into *35S::MLO15* and *mlo15-4.* Homozygous F3 seedlings were grown in FEP tubing as described above and imaged on a Bruker MuVi SPIM light sheet microscope with the same objectives and set up as described above. Images were acquired with 10% 561 nm laser and a 580-627 BP filter with a line scan width of 50 pixels. A 50 ms exposure and 5 µm step size (for 3 steps) was used to image root hairs every 3 seconds for 12 minutes. The last 10 minutes of the image acquisition was used for further analysis to control for altered signal at the start due to laser illumination.

Sum intensity projections were generated in ImageJ and used for analysis. Images were cropped to only contain a single root hair and were converted from 16-bit to 8-bit. The TrackMate plugin in ImageJ was used to track the brightest signal in the root hair apex and mean fluorescence intensity was obtained for each frame (Tinevez *et al*., 2017). The brightest signal was detected as a 30 pixel-diameter region of interest (ROI) with the LoG detector in TrackMate and tracked in each frame. The mean fluorescence intensity was obtained from the ROI for each frame for calcium oscillation analysis. The background signal was subtracted for each frame individually. Intensity traces were normalized as ΔF/F_0_. We used the background subtracted R-GECO1 intensity in a 400 pixel^2^ window in the shank of the root hair as F_0_. Peak intensity extraction was performed in MATLAB with the *findpeaks* function. Fourier transform and power spectral density (PSD) analysis was performed in MATLAB as previously described (Candeo *et al*., 2017). The relevant MATLAB code can be found in link: https://github.com/SADDLab/Root_Hair_Calcium_Analysis_Toolkit.git.

### H2DCF-DA and Peroxy Orange 1 ROS staining in root hairs

Five day old vertically grown seedlings were stained using the protocol described in (Gayomba & Muday, 2020). For H2DCF-DA staining, seedlings were incubated in 25 µm CM-H2DCFDA (Thermo Fisher C6827) made from a 50 mM stock in DMSO and diluted in water for 4 minutes. Seedlings were mounted in water and imaged with a Zeiss LSM 880 upright confocal microscope with a 10x objective, 5% laser intensity, and 1 airy unit pinhole with GFP optics. For PO1 staining, seedlings were incubated in 50 µm PO1 (R&D Systems 4944) for 15 minutes in the dark. Seedlings were rinsed with water and mounted in water. Images were acquired with a 488 nm laser and 544-695 emission filter using 5% laser intensity and a pinhole size of 1 airy unit. For both stains, images of primary roots containing initiating and early growth stage root hairs were analyzed as 8-bit maximum intensity projections. For each root hair, mean fluorescence intensity was taken along a line from base to tip in ImageJ (Schindelin *et al*., 2012). Statistical comparison was done using a one-way ANOVA with a Tukey test in GraphPad Prism 10.

## Results

### faNTA is sufficient to complement *fer-4* root hair development

In previous studies, expression of the constitutively active faNTA chimeric protein under a synergid-specific promoter was sufficient to complement the *fer-1* fertility phenotype through restoring synergid [Ca^2+^]_cyt_ oscillations that occur in response to FER signaling upon pollen tube arrival (Ju *et al*., 2021; Gao *et al*., 2022). *fer* mutants also have problems maintaining root hair tip growth (Duan *et al*., 2010). [Ca^2+^]_cyt_ oscillations at the root hair tip are correlated with maintenance of root hair polarity and growth, but it is not clear if FER also regulates these [Ca^2+^]_cyt_ oscillations during root hair tip growth. Since faNTA is sufficient to bypass FER signaling in synergids and induce pollen tube bursting in *fer-1*, we hypothesized that increasing [Ca^2+^]_cyt_ in root hairs through faNTA expression would suppress the burst root hair phenotype of *fer* if [Ca^2+^]_cyt_ oscillations occur downstream of FER signaling as was seen in synergids. To test this hypothesis, faNTA was expressed in *fer* mutants under control of the constitutive Cauliflower Mosaic Virus 35S promoter (Odell *et al*., 1985) and the FER promoter to ensure that faNTA would be expressed in the same tissues as FER. Laser scanning confocal microscopy was used to observe 35S::faNTA-YFP localization in root hairs. In initiating, elongating, and mature root hairs faNTA is present at the plasma membrane (Fig S1a).

To test for complementation of the *fer* phenotype, early elongating root hairs on the primary root between 2-3 mm from the root tip were measured in Col-0, *fer-4*, and *35S::faNTA;fer-4* (Fig 1a). Compared to wild-type root hairs that had an average length of 271.8 µm, *fer-4* root hairs were significantly shorter with an average length of 83.4 µm. The reduced root hair length of *fer-4* was restored to 265.8 µm in *35S::faNTA;fer-4,* which is not significantly different from Col-0 (Fig 1b). The same complementation of root hair length was observed when *35S::faNTA* and *pFER::faNTA* were expressed in the *fer-1* allele in the Landsberg *erecta* (Ler) background (Fig S1b-c), indicating that promoter strength and genetic background did not play a role in the ability of faNTA to complement the *fer* root hair phenotype.

**Figure 1:**
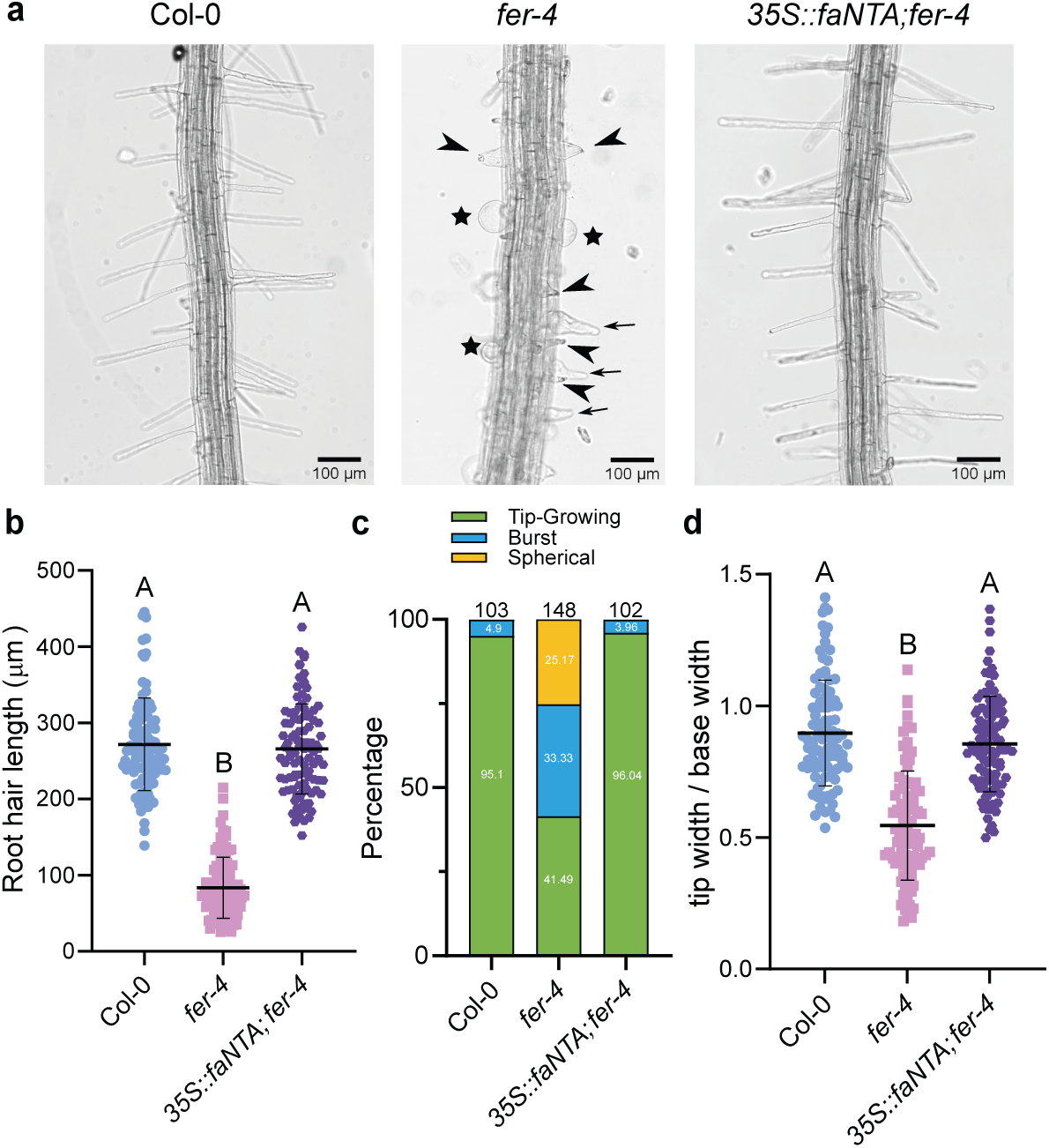
faNTA complements the abnormal root hair development of *fer-4*. A) Col-0, *fer-4*, and *35S::faNTA;fer-4* root hairs were imaged and phenotyped in a 1mm window 2-3mm from the primary root tip. For *fer-4,* stars indicate spherical root hairs, arrows indicate tip growing root hairs, and arrowheads indicate burst root hairs. Scale bar = 100 µm. B) The length of early elongating root hairs was measured for Col-0 (n=99), *fer-4* (n=101), and *35S::faNTA;fer-4* (n=99). C) The percentage of Col-0 (n=103), *fer-4* (n=148), and *35S::faNTA;fer-4* (n=102) root hairs that are tip-growing, burst, or spherical. D) The width of root hair tip divided by the width of root hair base was calculated for individual root hairs for Col-0 (n=100), *fer-4* (n=95), and *35S::faNTA;fer-4* (n=100). *fer-4* root hairs have wider root hair bases than tips. Multiple comparison analysis was done in GraphPad Prism10 as a one-way ANOVA with a Tukey test p<0.05.

In addition to the previously reported *fer* root hair bursting phenotype seen in 33% of root hairs, we observed two phenotypes that have not been previously described. The majority of *fer* root hairs in the early elongation zone were anisotropic and tip-growing but were cone-shaped with thicker bases than tips compared to wild-type root hairs. The remaining 25% of the root hairs were spherical and lacked polarized tip growth (Fig 1a, c-d). Spherical root hairs were never observed in Col-0 or *35S::faNTA;fer-4* root hairs. To determine whether *35S::faNTA* complements the altered tip-growing root hair shape in *fer-4*, the width of the root hair at the tip and base were measured and tip width/base width was calculated. Col-0 has a tip/base ratio of 0.89 and *fer-4* has a ratio of 0.55 (Fig 1d). *35S::faNTA;fer-4* root hairs have a tip/base ratio of 0.85, indicating that expression of faNTA is sufficient to complement root hair shape in a *fer-4* background (Fig 1d). The tip growing and spherical root hairs in *fer-4* are only found shortly after initiation and then eventually burst, which is likely why they have not been previously described.

### *fer* root hairs have altered Ca^2+^ oscillations that are complemented with faNTA

A tip-focused [Ca^2+^]_cyt_ gradient is important for maintaining tip integrity during root hair elongation (Monshausen *et al*., 2008). faNTA regulates Ca^2+^ influx when expressed in animal cells, and we hypothesized faNTA would also mediate Ca^2+^ influx in root hairs because it is plasma membrane localized (Fig S1a) (Gao *et al*., 2022). The loss of tip integrity in *fer-4* root hairs has been attributed to a reduction in ROS, but Ca^2+^ dynamics have not been explored in *fer-4* root hairs (Duan *et al*., 2010). We predicted that expression of faNTA in *fer-4* might lead to a larger influx of Ca^2+^ during oscillations that could contribute to root hair elongation and tip maintenance. To observe Ca^2+^ oscillations in wild-type, *fer-4,* and *35S::faNTA-YFP;fer-4* root hairs, the intensiometric cytoplasmic Ca^2+^ sensor R-GECO1 (Keinath *et al*., 2015) was crossed into these lines. Light sheet fluorescence microscopy (LSFM) allows seedlings to grow vertically in tubes that are then imaged vertically, reducing the amount of sample handling and more closely mimicking natural growing conditions. LSFM was used to image early elongating root hairs every 3 seconds for 10 minutes and R-GECO1 intensity at the root tip was measured, similar to the LSFM imaging and analysis done with Yellow Cameleon 3.6 in (Candeo *et al*., 2017). R-GECO1 intensity at the root hair apex was normalized to R-GECO1 intensity in the shank. Normalized intensity traces show that tip-growing *fer-4* root hairs have erratic oscillations that occur less frequently and with a larger variation in amplitude compared to Col-0 root hairs that oscillate with consistent frequency and amplitude over the imaging period (Fig 2a, additional traces S2, movie S1-2). In contrast to the tip-growing *fer-4* root hairs, spherical *fer-4* root hairs had [Ca^2+^]_cyt_ oscillations that occurred at very low amplitude and frequency so only tip-growing root hairs were used for further analysis. Normalized intensity traces of *35S::faNTA;fer-4* root hairs show that 35S::faNTA restores normal oscillation timing and amplitude to *fer-4* (Fig 2a, additional traces S2, movie S3).

**Figure 2:**
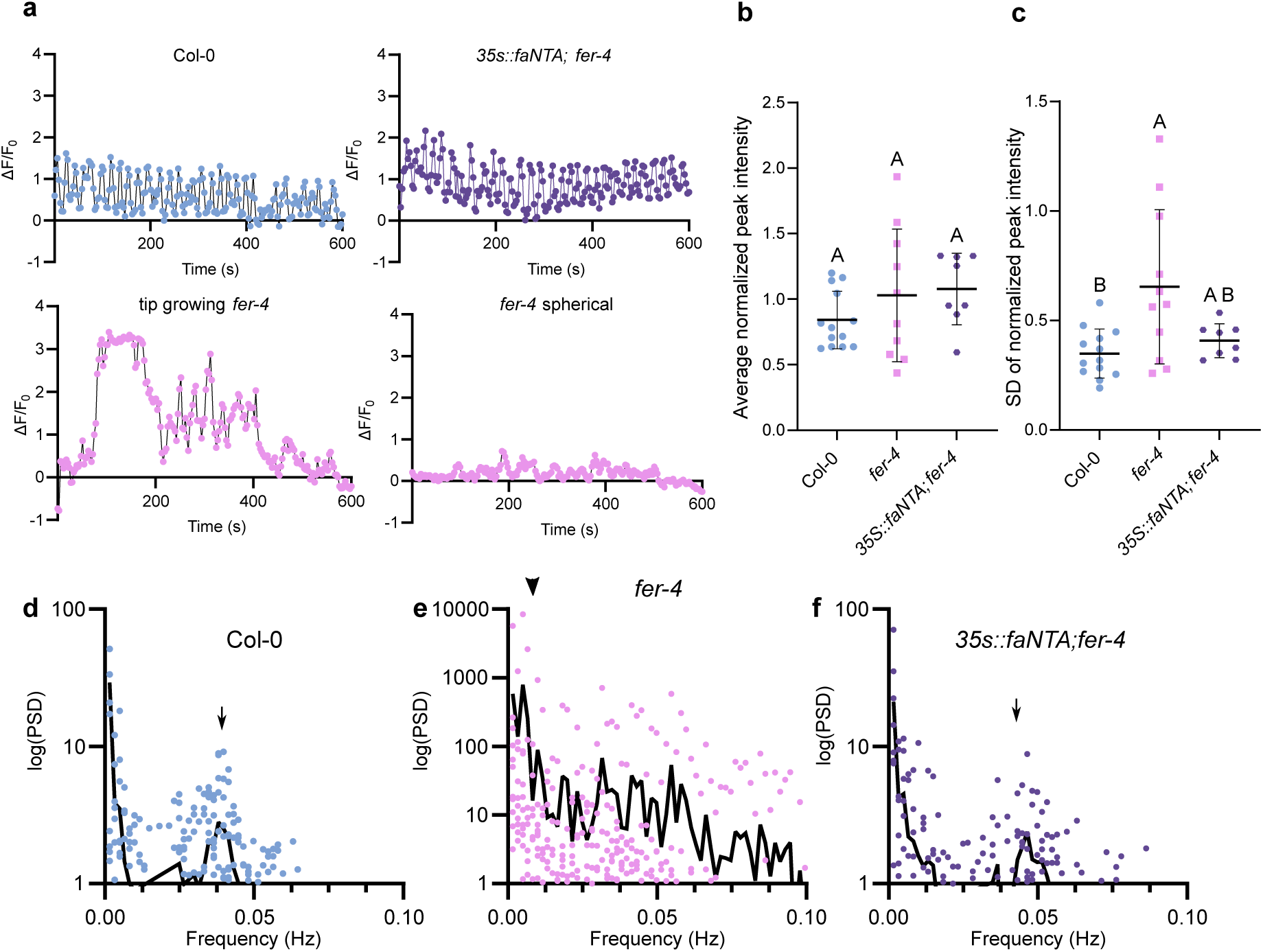
*fer-4* root hairs have Ca^2+^ oscillations of erratic amplitude and altered frequency. A) Normalized R-GECO1 intensity traces for Col-0, *35S::faNTA;fer-4*, tip-growing *fer-4*, and spherical *fer-4* root hairs. Additional traces can be found in Supplemental Figure 2. B-C) Normalized intensity at each peak was extracted for individual root hairs and the average intensity (B) or standard deviation (SD) from the average intensity (C) plotted for Col-0 (n=12), *fer-4* (n=11), *35S::faNTA;fer-4* (n=8). Multiple comparison was done in GraphPad Prism10 as a one-way ANOVA with a Tukey test p<0.05. D-I) Fourier transform and power spectral density (PSD) analysis results of R-GECO1 normalized signal traces (dots) and mean frequencies (black line) for Col-0 (D, G n=12), tip-growing *fer-4* (E,H n=11), and *35S::faNTA;fer-4* (F,I n=8). The lower PSD range is expanded in G-I to better show the variation in higher frequency oscillations. Arrows indicate the major high frequency peak for Col-0 and *35S::faNTA;fer-4* and the arrowhead indicates the major low frequency peak for *fer-4*.

To quantify the differences in [Ca^2+^]_cyt_ oscillation amplitude between Col-0, tip growing *fer-4*, and *35S::faNTA;fer-4* root hairs, the normalized intensity value for each oscillation peak was extracted for individual root hairs. The average normalized peak intensity is not different between lines (Fig 2b), likely because *fer-4* oscillations occur at amplitudes both higher and lower than Col-0. The standard deviation between normalized peak intensities were calculated for individual root hairs and plotted to better represent the range of amplitudes in *fer-4* tip growing root hairs (Fig 2c). The average standard deviation in peak intensity is higher in *fer-4* compared to the wild type and the average standard deviation for *35S::faNTA;fer-4* is not significantly different from both Col-0 and *fer-4* (Fig 2c). Within *fer-4,* standard deviations vary with some root hairs having similar values as Col-0 with some root hairs having much higher standard deviation (Fig 2c). This data along with the normalized intensity traces show that *fer-4* tip-growing root hairs have erratic [Ca^2+^]_cyt_ oscillations with altered amplitude.

*fer-4* root hairs appear to have fewer [Ca^2+^]_cyt_ oscillations than Col-0 in the normalized intensity traces (Fig 2a, S2). To quantify changes in [Ca^2+^]_cyt_ oscillation frequency, normalized R-GECO1 intensity data was Fourier transformed and the power spectral density calculated as described in (Candeo *et al*., 2017) to determine the interval of [Ca^2+^]_cyt_ oscillations in Col-0, *fer-4*, and *35S::faNTA;fer-4* root hairs. Col-0 root hairs have a single peak at 0.038 Hz, which corresponds to oscillations occurring every 26.3 seconds (Fig 2d, g). This result is consistent with the 0.036 Hz frequency reported for Col-0 root hairs expressing the ratiometric [Ca^2+^]_cyt_ reporter Yellow Cameleon 3.6 (YC3.6) imaged with light sheet microscopy and the 0.036-0.038 Hz frequency reported for Col-0 root hairs expressing R-GECO1 with confocal and RootChip imaging (Candeo *et al*., 2017; Brost *et al*., 2019). *fer-4* root hairs have a main low frequency peak at 0.0049 Hz, which corresponds to an oscillation every 204 seconds (Fig 2e, h). *fer-4* root hairs do not have a single high frequency peak, instead peaks occur from 0.0016-0.059 Hz (Fig 2e, h). The variability in frequency of Ca^2+^ oscillations suggests that one function of FER is to regulate Ca^2+^ influx in root hairs so oscillations occur at regular intervals. *35S::faNTA;fer-4* root hairs have a single peak at 0.046 Hz and oscillate every 21.7 seconds, which is a faster oscillatory pattern than Col-0 (26.3 seconds) (Fig 2f, i). Although it was known that FER regulates Ca^2+^ oscillations in synergids during pollen tube reception, these results demonstrate a new role for FER as a regulator of Ca^2+^ oscillations during root hair elongation. faNTA-mediated Ca^2+^ influx is able to complement both the altered amplitude and oscillation patterns in *fer-4* root hairs which are sufficient to restore normal tip growth to *fer* root hairs.

### MLO15 contributes to root hair tip growth

The ability of faNTA to complement *fer* root hair tip growth phenotypes suggests that MLOs may be an important downstream component of FER signaling in growing root hairs. Since faNTA is a chimeric protein and NTA is not expressed in root hairs, we utilized single-cell root transcriptomics data to identify candidate MLO genes expressed during root hair development (Ryu *et al*., 2019). MLO15 was the most highly expressed MLO in growing root hairs (Fig S3). Consistent with the single-cell RNA-seq data, plants expressing *pMLO15::MLO15-GFP* had higher GFP signal in root hairs compared to non-hair epidermal cells (Fig 3a).

**Figure 3:**
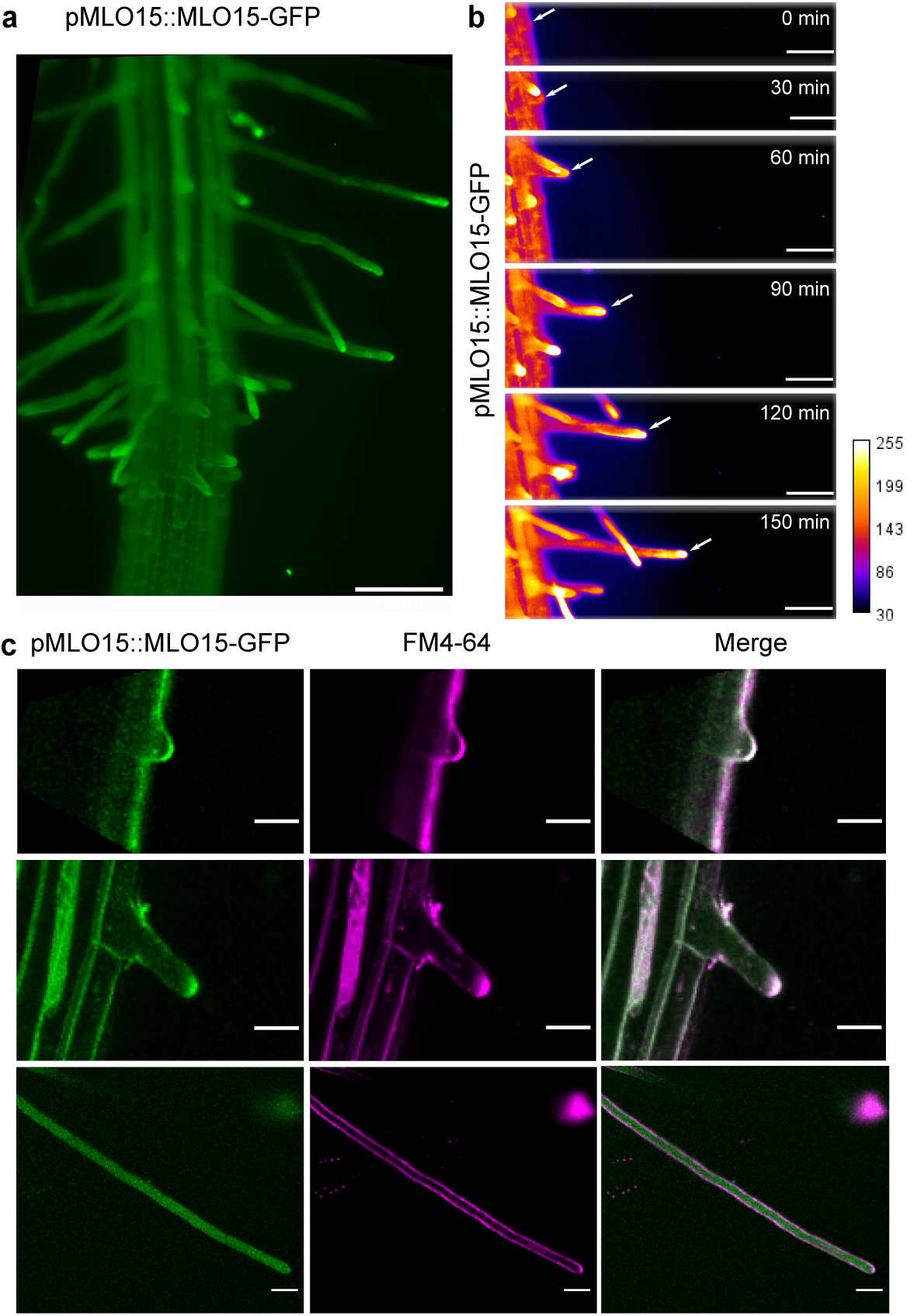
MLO15 is expressed in root hairs and its subcellular localization changes during root hair development. A) pMLO15::MLO15-GFP signal is expressed higher in root hair epidermal cells than non-hair epidermal cells. Scale bar = 100 µm. B) Light sheet microscopy was used to image MLO15-GFP during root hair initiation and elongation. During root hair initiation, MLO15-GFP signal is weak (0 min) and after rapid elongation begins, MLO15-GFP accumulates at the root apex. Images were pseudocolored to show MLO15-GFP intensity with white indicating the highest fluorescence. Scale bar = 50 µm. C) Laser scanning confocal microscopy was used to image MLO15-GFP in early, elongating, and mature root hairs. FM4-64 staining was used to distinguish the plasma membrane and the root apex. In the root hair bulge, MLO15-GFP accumulates at the plasma membrane. During elongation, MLO15-GFP signal is visible at the plasma membrane and also at the root apex. Mature root hairs show MLO15-GFP signal diffuse throughout the root hair. Scale bar = 20 µm.

To determine the root hair growth stages where MLO15 is expressed, LSFM was used to image *pMLO15::MLO15-GFP* roots every five minutes for four hours. This analysis revealed that MLO15-GFP signal is low and shows uniform signal in initiating root hairs and accumulates at the root tip during active root hair elongation (Fig 3b, movie S4). Confocal microscopy was used to determine the subcellular localization of MLO15-GFP at different stages of root hair development. In early elongating root hairs, MLO15-GFP is present at the plasma membrane of the root hair bulge (Fig 3c). When root hairs are actively elongating, MLO15-GFP localizes at the plasma membrane along the entire root hair, but also co-localizes with FM4-64 at the root tip (Fig 3c). FM4-64 labels the root hair apical zone vesicles along with the plasma membrane (Ovecka *et al*., 2005). In mature root hairs that have stopped growing, MLO15-GFP signal is found in punctate compartments evenly distributed throughout the cytosol and does not appear plasma membrane localized (Fig 3c). The co-localization of MLO15-GFP and FM4-64 in growing root hairs observed with confocal microscopy is consistent with the light sheet imaging data. MLO15 accumulates at the root hair tip and plasma membrane during elongation and is internalized in mature root hairs, suggesting that MLO15 is involved in root hair growth, but is not necessary for maintenance of mature root hairs.

Since root hairs are tip-growing cells, we hypothesized that MLO15 could be necessary for maintaining tip integrity or tip growth as these are both processes regulated by FER signaling. If this is true, then we expected that *mlo15* mutants would have similar root hair phenotypes to *fer* mutants. First, the length of early elongating root hairs was measured for Col-0, *mlo15-4*, and the complemented line *pMLO15::MLO15-GFP*;*mlo15-4.* Compared to Col-0, which had an average root hair length of 271.8 µm, *mlo15-4* had shorter root hairs that had an average length of 149.2 µm (Fig 4a-b). The decreased root hair length of *mlo15-4* was restored to wild-type levels by expressing *pMLO15::MLO15-GFP* and the complemented line had an average length of 295.1 µm (Fig 4a-b). To better characterize the reduced root hair length of *mlo15-4*, primary roots of 5-day old Col-0, *mlo15-4*, and the complemented line *pMLO15::MLO15-GFP;mlo15-4* were imaged and hairs along the entirety of the primary root were measured. When root hair length was plotted according to the position of that root hair on the primary root, all three genotypes have root hairs that increase in length as they mature. However, *mlo15-4* root hairs only reach half the length of Col-0 root hairs, with maximum length of approximately 400 µm compared to 800 µm for Col-0 and 900 µm for the complemented line (Fig 4e). In *fer-4* mutants, 33.33% of root hairs burst (Fig 1c). The *mlo15-4* mutant has on average 21% root hair bursting compared to the 4% root hair bursting in Col-0 and 8% in the complemented *mlo15-4* line (Fig 4d). However, in *mlo15-4* root hair bursting phenotype is not consistent among seedlings imaged. This suggests that *mlo15-4* roots may be more sensitive to osmotic changes caused from transfer from solid plates to water on slides leading to the increase in burst root hairs. We also examined root hair shape in the *mlo15* mutant to see if they are similar to the cone-shaped *fer-4* root hairs. *mlo15-4* root hairs have a slight decrease in the tip width/base width ratio compared to the wild-type. This phenotype is complemented with *pMLO15::MLO15-GFP* (Fig 4c). Overall, the differences in root hair architecture and shape observed in *mlo15-4* are similar but more subtle than the ones observed for *fer-4* (Fig 1a-d), indicating that other Ca^2+^ regulators may play redundant roles downstream of FER.

**Figure 4:**
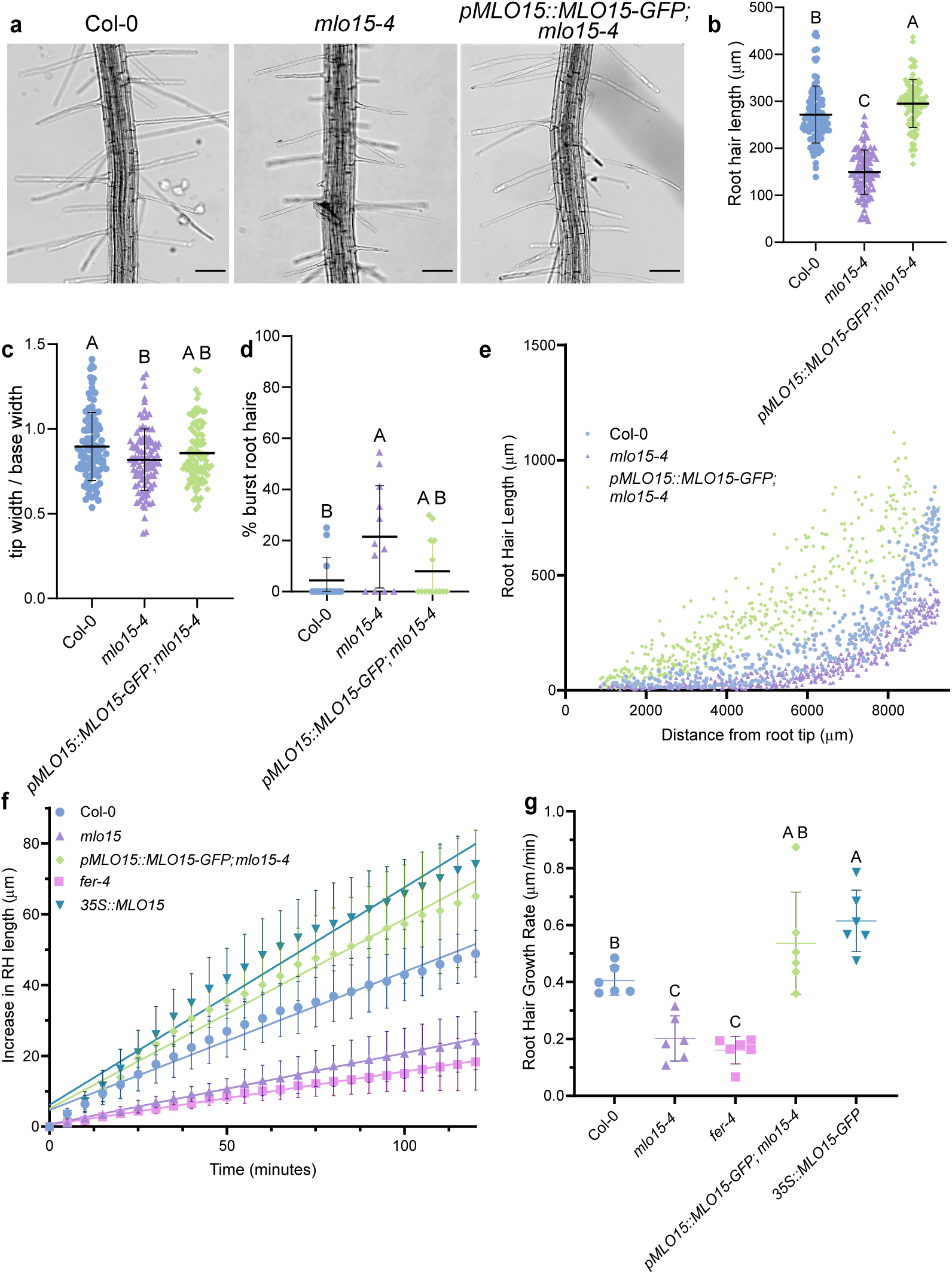
MLO15 contributes to root hair elongation. A-D) Col-0, *mlo15-4*, and *pMLO15::MLO15-GFP;mlo15-4* root hairs were imaged and phenotyped in a 1 mm window 2-3 mm from the primary root tip (A). Root hair length, tip width/base width, and bursting was quantified for Col-0 (n= 13 seedlings or 100 root hairs), *mlo15-4* (n=12 seedlings or 98 root hairs), and *pMLO15::MLO15-GFP;mlo15-4* (n= 13 seedlings or 101 root hairs) (B-D). In B-C, each data point represents a single root hair, while in D each data point represents the average of a single root. In the early root hair development region analyzed, *mlo15-4* root hairs are shorter (B), have a slightly lower tip width to base width ratio (C), and have a larger percentage of burst root hairs (D) compared to Col-0 that is complemented with pMLO15::MLO15-GFP. E) *mlo15-4* root hairs (n=465) along the entire primary root have reduced length compared to Col-0 (n=405) and *pMLO15::MLO15-GFP;mlo15-4* (n=453). Each data point represents a single root hair. F-G) MLO15 regulates root hair growth rate. Root hair length was measured every 5 minutes for 2 hours for individual root hairs Col-0 (n=6), *fer-4* (n=6), *mlo15-4* (n=6), *pMLO15::MLO15-GFP;mlo15-4* (n=5), and *35S::MLO15* (n=6). Averages are shown in (F) with standard errors. Root hair growth rate was calculated for individual root hairs by calculating the slope of the regression lines in F. *mlo15-4* and *fer-4* root hairs elongate at half the rate of the wild-type (G). Scale bar = 100 µm. Linear regression and statistical comparisons were done in GraphPad Prism 10 with a one-way ANOVA with a Tukey test p<0.05.

Since the *mlo15-4* mutant had significantly shorter mature root hairs than wild-type roots, we hypothesized that the *mlo15-4* root hairs grow more slowly than wild-type root hairs. To directly measure root hair growth rates for Col-0, *fer-4, mlo15-4*, *35S::MLO15* and the complemented line, seeds were germinated and grown vertically on plates for 5 days then transferred to slides with liquid 1/10 MS and grown vertically overnight before imaging. Root hairs nearest to the tip were imaged at five-minute intervals for two hours and lengths were measured. The change in root hair length from the starting length was graphed over time and fit with a linear regression curve to estimate root hair growth rate (Fig 4f). Consistent with our hypothesis, over the two-hour period, wild-type Col-0 root hairs elongated at a rate of 0.41 µm/min while *fer-4* root hairs elongated at 0.16 µm/min (half of the root hairs burst during the imaging window and were not measured). The *mlo15-4* mutants had a growth rate comparable to *fer-4* at 0.20 µm/min, but did not exhibit the high rate of root hair bursting of *fer-4* (Fig 4g). The complemented *mlo15-4* line and *35S::MLO15* lines elongated even faster than the wild type at rates of 0.54 µm/min and 0.61 µm/min, respectively (Fig 4g). These genotypes also had the greatest variation in growth rate between seedlings of the same genotype.

The root hair growth rates in all genotypes tested were not linear over the entirety of the imaging window, with the first hour having a faster growth rate than the second hour (Fig 4f). This might be attributed to seedling response to altered environmental conditions during imaging and being imaged horizontally, not vertically.

### MLO15 regulates Ca^2+^ during root hair growth

Since *mlo15-4* mutants have a similar root hair growth phenotype to *fer-4*, we hypothesized that the reduced root hair elongation rate in *mlo15-4* would also be correlated with reduced [Ca^2+^]_cyt_ levels. In HEK293 cells, MLO15 was shown to be capable of conducting Ca^2+^ when co-expressed with upstream signaling components (Gao *et al*., 2023). However, MLO15 has not been shown to regulate [Ca^2+^]_cyt_ levels in plant cells. To test whether the abnormal root hair development of *mlo15-4* is due to reduced [Ca^2+^]_cyt_, the Ca^2+^ sensor R-GECO1 in Col-0 was crossed into *mlo15-4* and *35S::MLO15-GFP*. LSFM was used to observe R-GECO1 intensity in young root hairs that have started to elongate. Images were acquired every 3 seconds for 10 minutes. If MLO15 functions as a Ca^2+^ channel in root hairs, we might observe reduced amplitude or altered frequency in *mlo15-4* root hairs. Normalized intensity traces show that *mlo15-4* root hairs have oscillations that occur at a reduced amplitude compared to Col-0 and *35S::MLO15-GFP* root hairs have oscillations that appear similar to Col-0 (Fig 5a, additional traces S4, movies S2, 5, 6). Normalized peak intensity was extracted for individual root hairs and the average peak intensity was compared (Fig 5b). Compared to Col-0, *mlo15-4* has a significantly reduced average peak intensity and *35S::MLO15-GFP* has a similar average peak intensity (Fig 5b). This data shows that *mlo15-4* root hairs have [Ca^2+^]_cyt_ oscillations that occur at a reduced amplitude.

**Figure 5:**
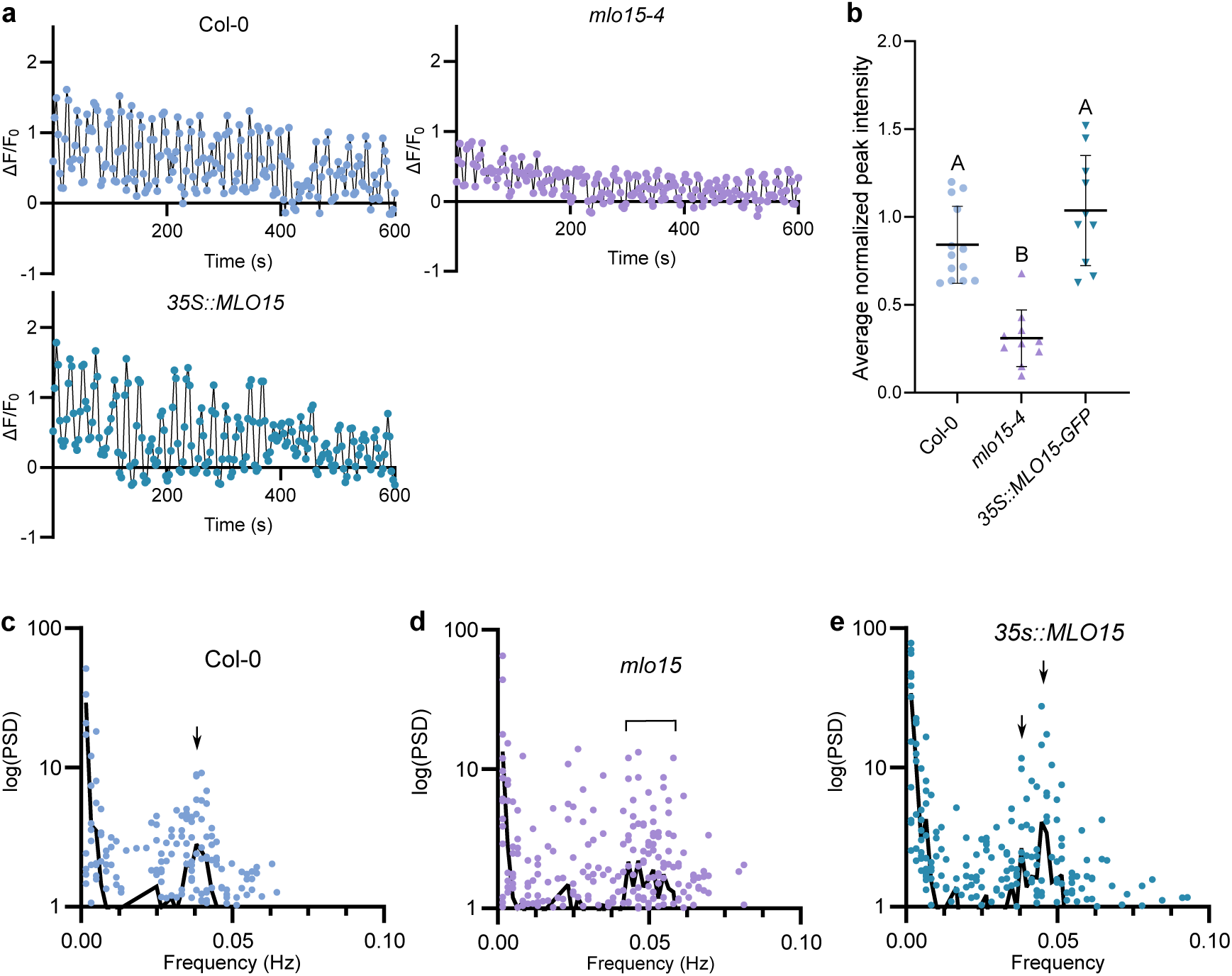
*mlo15-4* root hairs have Ca^2+^ oscillations of reduced amplitude and altered frequency. A) Representative normalized R-GECO1 intensity traces for Col-0, *mlo15-4,* and *35S::MLO15-GFP* root hairs imaged every 3 seconds for 10 minutes. Additional traces can be found in Supplemental Figure 4. B) Normalized peak intensity was extracted for individual root hairs and averages were plotted for Col-0 (n=12), *mlo15-4* (n=10), and *35S::MLO15-GFP* (n=10). Multiple comparison was done with a one-way ANOVA with Tukey test in GraphPad Prism10 p<0.05. C-H) Fourier transformation results of R-GECO1 normalized signal traces (dots) and mean frequencies (black line) for Col-0 (C,F n=12), *mlo15-4* (D,G n=17), and *35S::MLO15-GFP* (E,H n=12). The lower PSD range is expanded in F-H to better show the variation in higher frequency oscillations. Arrows indicate main high frequency peaks for Col-0 and *35S::MLO15*, bracket indicates high frequency range for *mlo15-4.* The Col-0 data shown in panels a, b, c, and f are duplicated from the data shown in Figure 2 for easier comparison.

Altered Ca^2+^ oscillation frequency in root hairs was observed in *fer-4* (Fig 2) and reported in two double mutants of CNGC-type Ca^2+^ channels, *cngc6;14* and *cngc9;14* (Brost *et al*., 2019). As previously described in Figure 2d and 2g, Col-0 root hairs had a single peak at 0.038 Hz, which corresponds to oscillations occurring every 26.3 seconds. *mlo15-4* root hairs show a loss of regular oscillations and have an extended peak from 0.04 to 0.06 Hz that suggests these root hairs oscillate more frequently than Col-0 with oscillations every 17-25 seconds (Fig 5d, g). *fer-4* root hairs also had disrupted oscillation patterning with a large low frequency peak at 0.0049 Hz and an extended peak from 0.0016-0.059 Hz (Figure 2De and 2h). Compared to *fer-4* oscillations, *mlo15-4* oscillations do not show a low frequency peak and the PSD distribution is less varied. *35S::MLO15-GFP* root hairs have two high frequency peaks:, one at 0.038 Hz, which is the same as Col-0, and one at 0.044 Hz, which corresponds to a 23 second oscillation (Fig 5e, h). Together, these data support a function for MLO15 as a regulator of Ca^2+^ influx in root hairs that contributes to normal [Ca^2+^]_cyt_ oscillations required for root hair growth.

### MLO-signaling promotes ROS production in root hairs

The loss of tip integrity in *fer-4* root hairs has previously been attributed to a reduction in ROS (Duan *et al*., 2010). FER promotes ROS production through ROP-GEF signaling that activates RBOHC (Duan *et al*., 2010). ROS signaling has been proposed to regulate Ca^2+^ channels, and Ca^2+^ has also been linked to regulation of RBOH activity (Véry & Davies, 2000; Takeda *et al*., 2008; Zhang *et al*., 2018). We have shown that faNTA can bypass FER signaling in root hairs to restore normal [Ca^2+^]_cyt_ levels to *fer-4*. The next question is whether faNTA can also restore ROS levels in a FER-independent manner.

To determine whether faNTA complementation of *fer-4* also complements the reduction in root hair ROS levels, the non-specific cell-permeable ROS detector H2DCF-DA was used. ROS levels in Col-0, *fer-4*, and *35S::faNTA;fer-4* root hairs were quantified (Fig 6a-b). *fer-4* root hairs had a third of the H2DCF-DA signal as Col-0, while *35S::faNTA;fer-4* was not significantly different (Fig 6b). This demonstrates that faNTA is sufficient to increase ROS levels in root hairs independent of FER signaling. H2DCF-DA is a general ROS dye that has previously been used to characterize the reduced ROS levels in *fer* mutants (Duan *et al*., 2010; Kim *et al*., 2021). To determine whether H_2_O_2_ is increased, which would be consistent with faNTA activating RBOHC, the dye Peroxy Orange 1 (PO1) that is cell-permeable and specific for H_2_O_2_ was used (Fig 6c-d). Consistent with the H2DCF-DA staining, *fer-4* had a quarter of the PO1 fluorescence compared to Col-0 and *35S::faNTA;fer-4* (Fig 6d). This staining was repeated on the *fer-1* allele in the Ler background and faNTA was sufficient to restore ROS levels in *fer-1* with both H2DCF-DA and PO1 staining (Fig S5). These data suggest that faNTA promotes ROS production in root hairs independently of FER signaling.

**Figure 6:**
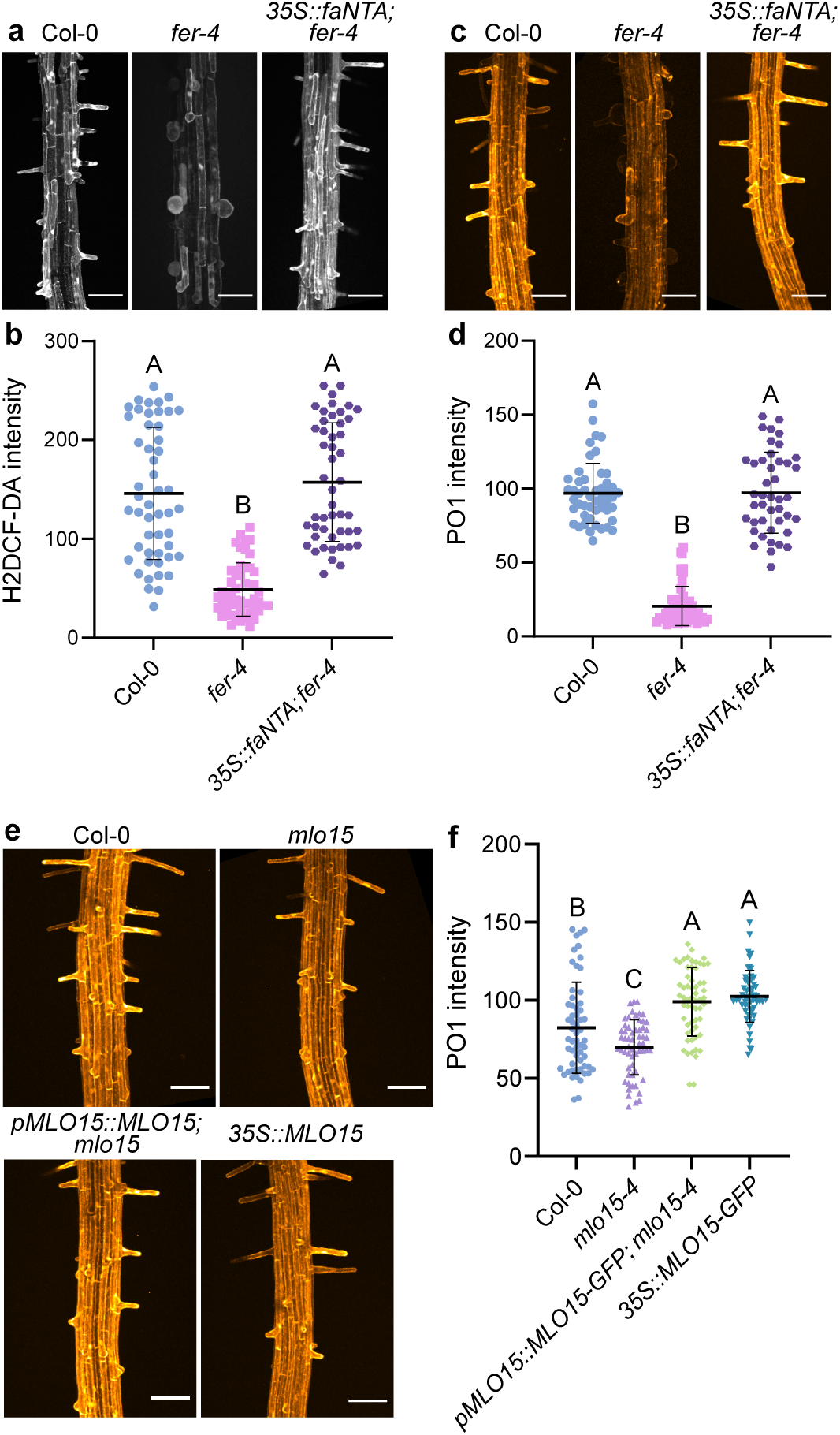
faNTA and MLO15 positively regulate ROS production in root hairs. A-B) The general ROS indicator H2DCF-DA was used to compare ROS levels in Col-0 (n=50), *fer-4* (n=48), and *35S::faNTA;fer-4* (n=48) root hairs. C-D) PO1 staining was used to quantify H_2_O_2_ levels in Col-0 (n=49), *fer-4* (n=54, and *35S::faNTA;fer-4* (n=43) root hairs E-F) PO1 staining was used to quantify H_2_O_2_ levels in Col-0 (n=58), *mlo15-4* (n=58), *pMLO15::MLO15-GFP;mlo15-4* (n=59), and *35S::MLO15-GFP* (n=57) root hairs. Fluorescence intensity was compared in GraphPad Prism 10 with a one-way ANOVA with Tukey test p<0.05. Scale bar = 100 µm.

Since the constitutively activated MLO, faNTA, complements ROS in *fer-4* root hairs, we hypothesized that the root hair-expressed MLO15 also has a role regulating ROS production in root hairs. We expected that if MLO15 positively regulates ROS levels, *mlo15-4* root hairs would have reduced ROS and the *35S::MLO15-GFP* root hairs might have increased ROS. PO1 staining was done on wild-type Col-0, *mlo15-4*, *pMLO15::MLO15-GFP*;*mlo15-4*, and *35S::MLO15-GFP* (Fig 6e-f). Compared to Col-0, *mlo15-4* has a slight, but significant reduction in PO1 signal and the complemented line and *35S::MLO15-GFP* root hairs have increased signal, indicating that MLO15 may have a role in regulating ROS production in growing root hairs (Fig 6f).

## Discussion

Root hair tip growth involves coordinated Ca^2+^ and ROS signaling to promote growth while maintaining tip integrity. In this study, we took advantage of the constitutively active faNTA chimeric protein to identify a new role for the FER/MLO signaling module in regulating [Ca^2+^]_cyt_ oscillations in growing root hairs. We previously demonstrated that faNTA is sufficient to bypass *fer* signaling in synergids. Interestingly, faNTA is also able to bypass FER signaling in root hairs, complementing not only the previously reported *fer-4* root hair bursting phenotype, but also the complete loss of polarity in spherical root hairs. This complementation correlated with the restoration of tip-focused [Ca^2+^]_cyt_ oscillations that are disrupted in *fer* mutants. The faNTA complementation data suggested that MLO proteins might be involved in regulating [Ca^2+^]_cyt_ status in growing root hairs. We showed that MLO15 is the MLO family member involved in regulating root hair tip growth. The amplitude of [Ca^2+^]_cyt_ oscillations are reduced in *mlo15-4* and oscillations occur at a higher frequency than in the wild type, leading to slower growth and mild *fer*-like phenotypes. We also link the FER/MLO module to ROS accumulation by showing that faNTA is sufficient to restore ROS levels in *fer-4* root hairs and that ROS levels are decreased in *mlo15* root hairs. We propose that one of the functions of the Ca^2+^ influx mediated by MLOs in root hairs is to promote ROS production.

### FER and MLO15 contribute to the regulation of [Ca^2+^]_cyt_ oscillation amplitude and timing in root hairs

Our study identified new roles for FER and MLO15 regulating [Ca^2+^]_cyt_ oscillations during root hair elongation. Both *fer-4* and *mlo15-4* root hairs have oscillations with altered amplitude and disrupted frequency. *mlo15-4* root hairs had a slight increase in root hair bursting compared to the wild-type, but do not have altered root hair initiation. This is consistent with our observation that MLO15 is not highly expressed during root hair initiation, accumulates at the root hair apex and plasma membrane during elongation, and is internalized at root hair maturity. The only other plasma membrane localized Ca^2+^ channels that have been reported to regulate root hair development are CNGC5, 6, 9, 14. Single and multiple mutants in the root hair CNGCs have similar phenotypes to *mlo15-4.* The shorter root hair length of *mlo15-4* is similar to the phenotype previously reported for *cngc6,9,14* and is consistent with pharmacological studies that suggest Ca^2+^ influx is not necessary for the formation of root hair bulges (Schiefelbein *et al*., 1992; Brost *et al*., 2019).

In root hairs, [Ca^2+^]_cyt_ and growth rate oscillations are correlated with [Ca^2+^]_cyt_ peaks lagging growth peaks by 5-7 seconds (Monshausen *et al*., 2008; Candeo *et al*., 2017). However, the relevance of the frequency of [Ca^2+^]_cyt_ oscillations in regulating root hair tip growth rates remains largely unknown. Analyses of calcium dynamics in *mlo15-4* and *cngc* mutants suggest that there is a not a linear relationship between oscillation frequency and growth rate. The frequency of [Ca^2+^]_cyt_ oscillations is shifted slightly lower in *cngc9,* while *cngc6,14* and *cngc9,14* root hairs have a loss of normal oscillations, similar to our observations in *fer-4.* The reduced amplitude and faster [Ca^2+^]_cyt_ oscillations of *mlo15-4* is similar to the *cngc14* phenotype where root hairs have faster oscillations that occur at roughly half the amplitude of Col-0 and have a slower growth rate (Zhang *et al*., 2017; Brost *et al*., 2019). One hypothesis for the reduced growth rate in *mlo15-4* and *cngc14* root hairs despite having more frequent [Ca^2+^]_cyt_ oscillations is that [Ca^2+^]_cyt_ needs to reach a certain threshold to activate downstream pathways. Further work can be done to dissect how [Ca^2+^]_cyt_ oscillation frequency and amplitude contribute to tip integrity and growth.

Previous studies have shown protein-protein interactions between other members of the MLO and CNGC families (Chen *et al*., 2012; Meng *et al*., 2020). Since the phenotypes of *mlo15* and the root hair expressed CNGC mutants are similar, it is possible that MLO15 interaction with CNGCs in root hairs regulates the amplitude and timing of [Ca^2+^]_cyt_ oscillations. Both CNGC and MLO channel activity are thought to be negatively regulated by CALMODULIN (CaM) binding at the C-terminus (Fischer *et al*., 2013; Fischer *et al*., 2017; Gao *et al*., 2022). Whether CaM binding functions to negatively regulate Ca^2+^ influx and maintain [Ca^2+^]_cyt_ oscillations in root hairs needs further investigation.

### [Ca^2+^]_cyt_ and ROS crosstalk during root hair development

[Ca^2+^]_cyt_ oscillations and a tip-focused ROS gradient function synergistically to promote tip growth of root hairs (Monshausen *et al*., 2009). One function of the ROS generated by RBOHC is to promote Ca^2+^ influx in root hair tips. The *rhd2/rbohc* mutant has root hairs with altered tip integrity that burst. *rhd2* root hairs also have lower [Ca^2+^]_cyt_ levels that can be increased with the addition of ROS (Foreman *et al*., 2003). The *rhd2* tip integrity phenotype is rescued and tip-focused [Ca^2+^]_cyt_ oscillations are restored when the media pH is increased to 6 (Monshausen *et al*., 2007). This suggests that one function of the ROS generated by RBOHC is to activate Ca^2+^ influx channels, but that RBOHC is not essential for formation of the [Ca^2+^]_cyt_ gradient. The abnormal root hair development of *fer-4* was previously attributed to a reduction of ROS (Duan *et al*., 2010). Our results confirm the reduced ROS levels in *fer-4* and also show *fer-4* root hairs have altered [Ca^2+^]_cyt_ oscillations with lower frequency and reduced amplitude. An outstanding question remains whether FER regulates [Ca^2+^]_cyt_ oscillations through direct regulation of Ca^2+^ channels or if the reduction in ROS in *fer-4* leads to altered regulation of Ca^2+^ channels and disrupted [Ca^2+^]_cyt_ oscillations.

Our finding that faNTA was able to complement ROS production in *fer* root hairs suggests that the increased [Ca^2+^]_cyt_ induces ROS production independently from the well-described FER/ROP-GEF/RBOHC signal transduction pathway described in (Duan *et al*., 2010). Ca^2+^ signaling can activate RBOHC to produce ROS in root hairs through multiple mechanisms. Ca^2+^-dependent phosphorylation of RBOHC through CALCINEURIN B-LIKE PROTEIN 1 (CBL1) and CBL-INTERACTING PROTEIN KINASE 26 (CIPK26) can activate RBOHC, but is not required for normal root hair growth (Zhang *et al*., 2018). Alternatively, RBOHC has two cytosolic EF-hand domains that can bind Ca^2+^. The EF-hand domains are necessary for RBOHC function in root hairs and when these EF-hands are mutated to abolish Ca^2+^ binding, RBOHC cannot complement the *rhd2* phenotype (Takeda *et al*., 2008). faNTA is able to conduct Ca^2+^ when expressed in animal cells and is sufficient to restore normal Ca^2+^ oscillations to *fer-4* (Gao *et al*., 2022). It is possible that the restoration of ROS in *fer-4* root hairs is through a mechanism where faNTA mediates Ca^2+^ influx that activates RBOHC through Ca^2+^ binding at the EF-hand motifs. More work needs to be done to dissect the mechanism of faNTA complementation of *fer-4* to determine if the complementation of ROS is in fact due to an increase in [Ca^2+^]_cyt_.

### FER functions as a regulator of Ca^2+^ signaling throughout the plant

Ca^2+^ signaling occurs in plant cells as a response to stress or other extracellular cues. In some cases, FER has been proposed to act as a sensor for cell wall perturbations and is required for the establishment of [Ca^2+^]_cyt_ oscillations. For example, *fer* roots have defective mechanosensing that is correlated with altered or abolished [Ca^2+^]_cyt_ signatures in response to touch (Shih *et al*., 2014). In salt stress experiments, the [Ca^2+^]_cyt_ signatures observed in primary roots recovering from salt stress-induced damage are absent in *fer-4* (Feng *et al*., 2018). During plant reproduction, FER is required for initiating [Ca^2+^]_cyt_ oscillations in synergids in response to signals from the arriving pollen tube. In contrast, our root hair data reveal that FER regulates the amplitude and frequency of [Ca^2+^]_cyt_ oscillations in root hairs, but is not necessary for the initiation of these oscillations. Nevertheless, it is clear that FER plays a role in ensuring that [Ca^2+^]_cyt_ oscillations are regulated properly for mounting appropriate cellular responses during growth and development and plant responses to the environment.

### MLOs as amplifiers of FER signaling

In plant reproduction, NORTIA (MLO7) was proposed to function as an amplifier of FER signaling in synergids to ensure pollen tube bursting and delivery of the sperm cells to the female gametes (Ju *et al*., 2021). As was seen in comparisons of *mlo15-4* and *fer-4* root hair phenotypes, *nta-1* plants have a less severe infertility phenotype than *fer* and are 30% infertile compared to *fer-1* plants that are 80% infertile (Escobar-Restrepo *et al*., 2007; Kessler *et al*., 2010). During pollen tube reception [Ca^2+^]_cyt_ oscillations occur in both the pollen tube and synergids (Ngo *et al*., 2014). *nta-1* synergids have [Ca^2+^]_cyt_ oscillations that occur at a reduced amplitude, while *fer-1* synergids have abolished oscillations (Ngo *et al*., 2014). Our lab has proposed a primary/secondary signaling mechanism in synergids during pollen tube reception where there is a threshold of pollen tube burst signal that needs to be reached for normal pollen tube reception to occur. FER signaling initiates a primary burst signal and also the activation of NTA, which then activates the secondary burst signal through amplification of [Ca^2+^]_cyt_ oscillations (Ju *et al*., 2021). faNTA induces pollen tube bursting in *fer-1* synergids that lack the primary burst signal, indicating that the secondary burst signal from faNTA is sufficient to reach the threshold needed to induce pollen tube bursting (Ju *et al*., 2021).

In this study, we showed that faNTA could also complement root hair growth phenotypes in *fer*, suggesting that a similar booster mechanism may be in play. It remains unclear how [Ca^2+^]_cyt_ oscillations are initiated in root hairs, but we now know that these oscillations are maintained through MLO and CNGC function (Zhang *et al*., 2017; Tan *et al*., 2020). It is possible that MLO15 in root hairs also functions as an amplifier of FER signaling in order to promote root hair integrity and growth in response to external signals, similar to how NTA functions to amplify the FER response signals from the arriving pollen tube. FER seems to be required for maintenance of root hair tip growth, with the majority of root hairs bursting or never establishing tip growth, while MLO15 seems to be a “helper” since *mlo15* root hairs grow slower but manage to reach maturity compared to the severe root hair bursting phenotype in *fer-4*.

Outside of pollen tube reception and root hair development, MLOs have been implicated in similar signaling processes as FER including powdery mildew susceptibility and root thigmotropism (Consonni *et al*., 2006; Chen *et al*., 2009; Kessler *et al*., 2010; Bidzinski *et al*., 2014; Shih *et al*., 2014; Darwish *et al*., 2022). In most instances, *fer* plants have a more severe phenotype than the *mlo* or multiple *mlos* need to be knocked out to display the same phenotype as *fer.* Further work can be done to determine whether there is a conserved role for MLOs functioning downstream of FER as amplifiers of FER signaling throughout the plant.

### How is MLO15 regulated?

We have used faNTA as a tool to investigate Ca^2+^ signaling in root hairs. faNTA is sufficient to restore normal [Ca^2+^]_cyt_ oscillations and ROS levels to *fer-4* root hairs, but it remains unclear whether FER is required for MLO15 activation in root hairs. Two modes of MLO activation have been described previously. (1) FER activates NTA (MLO7) by regulating its trafficking to the filiform apparatus in synergids and (2) the FER homologs ANX1/2 and BUPS1/2 activate MLO1/5/9/15 in pollen tubes through activating a cytoplasmic kinase intermediate MARIS (MRI) (Ju *et al*., 2021; Gao *et al*., 2022; Gao *et al*., 2023). MLO15 is unable to conduct Ca^2+^ when expressed by itself in HEK293 cells, but can mediate Ca^2+^ influx when co-expressed with upstream signaling genes (Gao *et al*., 2023). This suggests that MLO15 requires direct activation by another protein in order to function. It is possible that MLO15 activation in root hairs is similar to its activation in pollen tubes where FER activates MRI that then activates MLO15. A constitutively active *mri* mutant can complement *fer* root hair phenotypes, but MRI has not been implicated in Ca^2+^ signaling in root hairs and it remains unknown whether *mri* root hairs have altered [Ca^2+^]_cyt_ oscillations (Boisson-Dernier *et al*., 2015; Liao *et al*., 2016). Alternatively, FER has been shown to form signaling complexes with other CrRLKL1 proteins in other cell types (Galindo-Trigo *et al*., 2020; Liu *et al*., 2021). CAPS1/ERU is a CrRLKL1 protein that is also required for normal [Ca^2+^]_cyt_ oscillations in root hairs (Kwon *et al*., 2018). It is possible that MLO15 is a target of ERU or an ERU/FER complex. A third possibility is that the FER regulates MLO15 subcellular localization. MLO15 has different subcellular localization patterns in growing vs. mature root hairs which could be analogous to the FER/NTA system where NTA is retained in the Golgi until FER signaling is activated by the arriving pollen tubes, upon which it accumulates at the filiform apparatus, which is part of the plasma membrane of the synergid (Ju *et al*., 2021). A similar mechanism could be employed in root hairs, where FER signaling is on during active growth and leads to accumulation of MLO15 at the plasma membrane. When root hairs reach maturity and no longer need to grow, FER signaling would be turned down, inhibiting MLO15 accumulation in the plasma membrane. At this time, we also cannot rule out the possibility that MLO15 is regulated independently of FER in root hairs and FER instead activates other families of Ca^2+^ channels. Future work should focus on elucidating how FER regulates [Ca^2+^]_cyt_ oscillations in root hairs and to determine if this is through direct activation of MLO15.

## Acknowledgements

We thank Marilyn Vargas and Molly Kosiba for assisting with experiments, the Bindley Bioscience Center Imaging Facility for assistance with confocal microscopy, and the Purdue Plant Growth Facility assistance with plant cultivation. We thank the Purdue Office of the Vice President for Research for providing funding for the Bruker light sheet microscope. We also thank Dr. Leonor Boavida, Dr. Anjali Iyer-Pascuzzi, Dr. Tesfaye Mengiste, and Sowmiya Devi Venkatesan for helpful discussions.

## Author Contributions

STO, WZ, CJS and SAK conceived and designed the experiments. STO and WZ performed the experiments. STO, WZ, and SAK analyzed the data. STO and SAK wrote the manuscript, and all authors revised and approved the final manuscript.

## Data availability

Sequence data from this article can be found in GenBank/EMBL data libraries under accession numbers AT3G51550 (FER) and AT2G44110 (MLO15). The relevant MATLAB code can be found in link: https://github.com/SADDLab/Root_Hair_Calcium_Analysis_Toolkit.git. Any additional information required to reanalyze the data in this paper is available from the corresponding author upon request.

## Supporting information

**Supplemental Figure 1:**
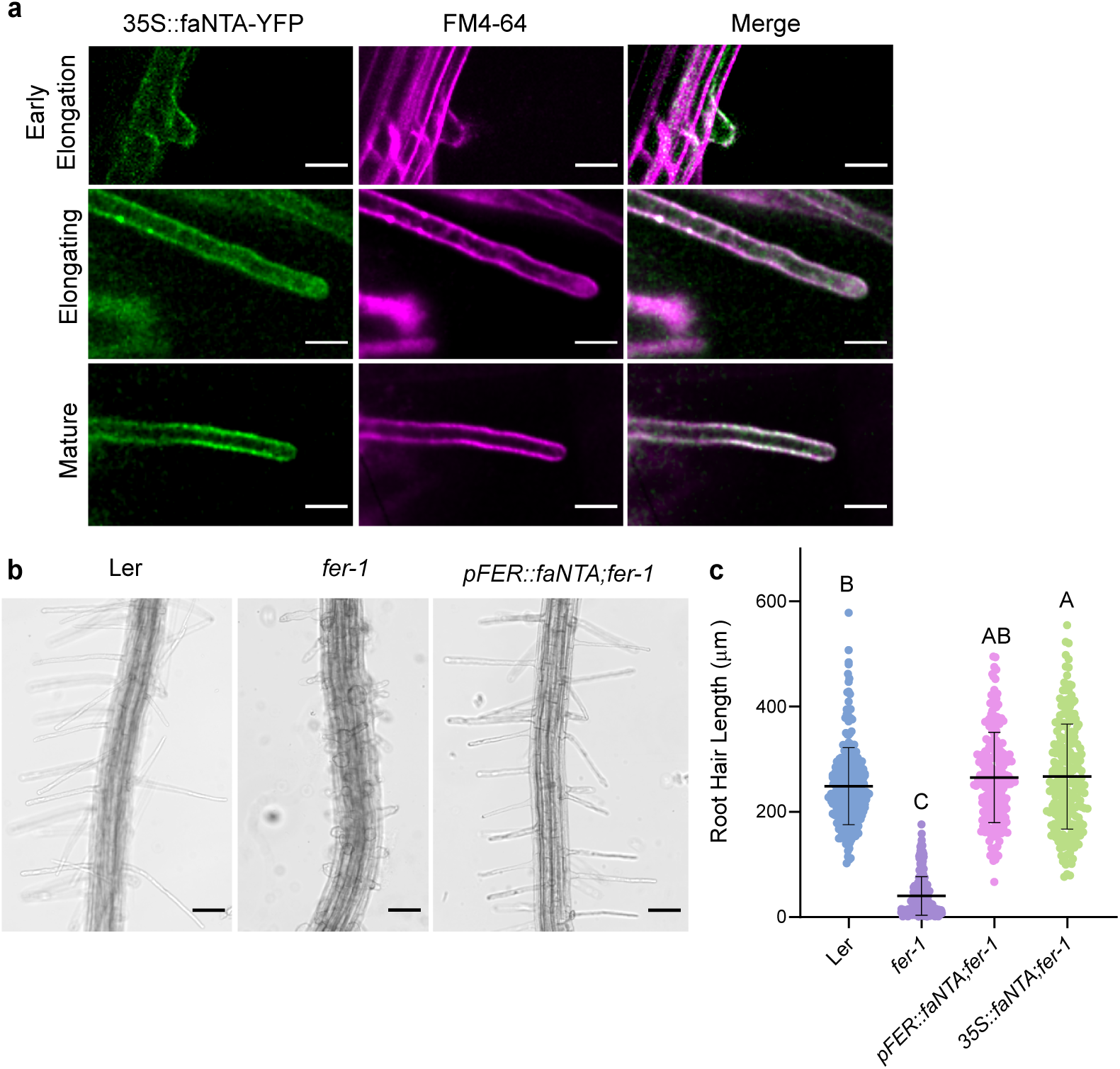
faNTA is plasma membrane localized and complements the abnormal root hair development of *fer-1*. A) 35S::faNTA-YFP in *fer-4* subcellular localization during root hair growth was observed with laser scanning confocal microscopy. faNTA-YFP co-localizes with FM4-64 at the plasma membrane at all growth stages and at the root hair apex in elongating root hairs. Scale bar = 20 µm. B) Ler, *fer-1*, and *<u>pFER::faNTA;</u>fer-1* early elongating root hairs images. Scale bar = 100 µm. C) The length of early elongating root hairs was measured for Ler (n=281), *fer-1* (n=281), *pFER::faNTA;fer-1* (n=217), and *35S::faNTA;fer-1* (n=219). Multiple comparison was done in GraphPad Prism 10 with a one-way ANOVA with a Tukey test p<0.05.

**Supplemental Figure 2:**
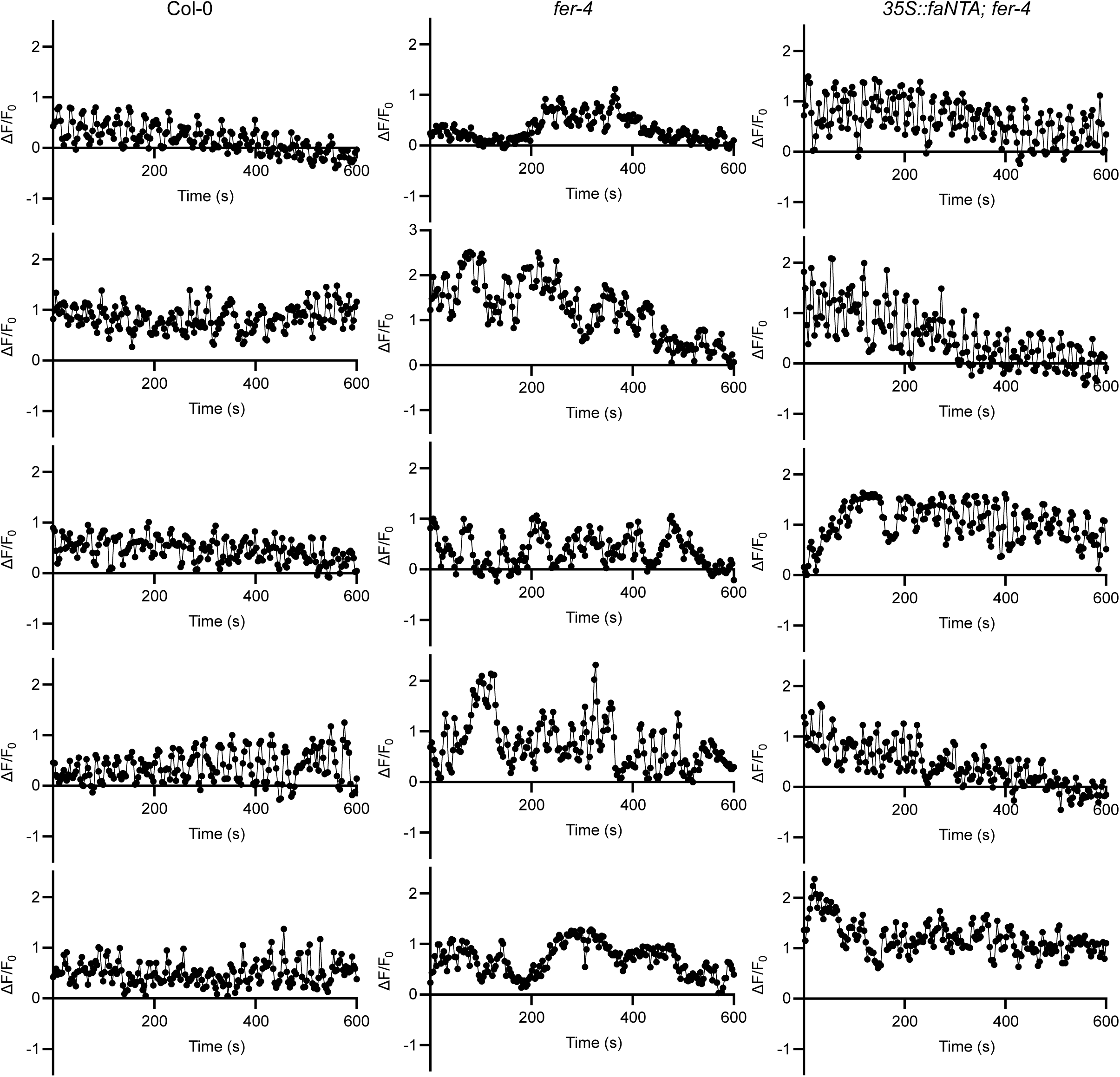
Additional normalized intensity traces of R-GECO1 intensity over 10 minutes in Col-0, *fer-4*, and *35S::faNTA;fer-4*.

**Supplemental Figure 3:**
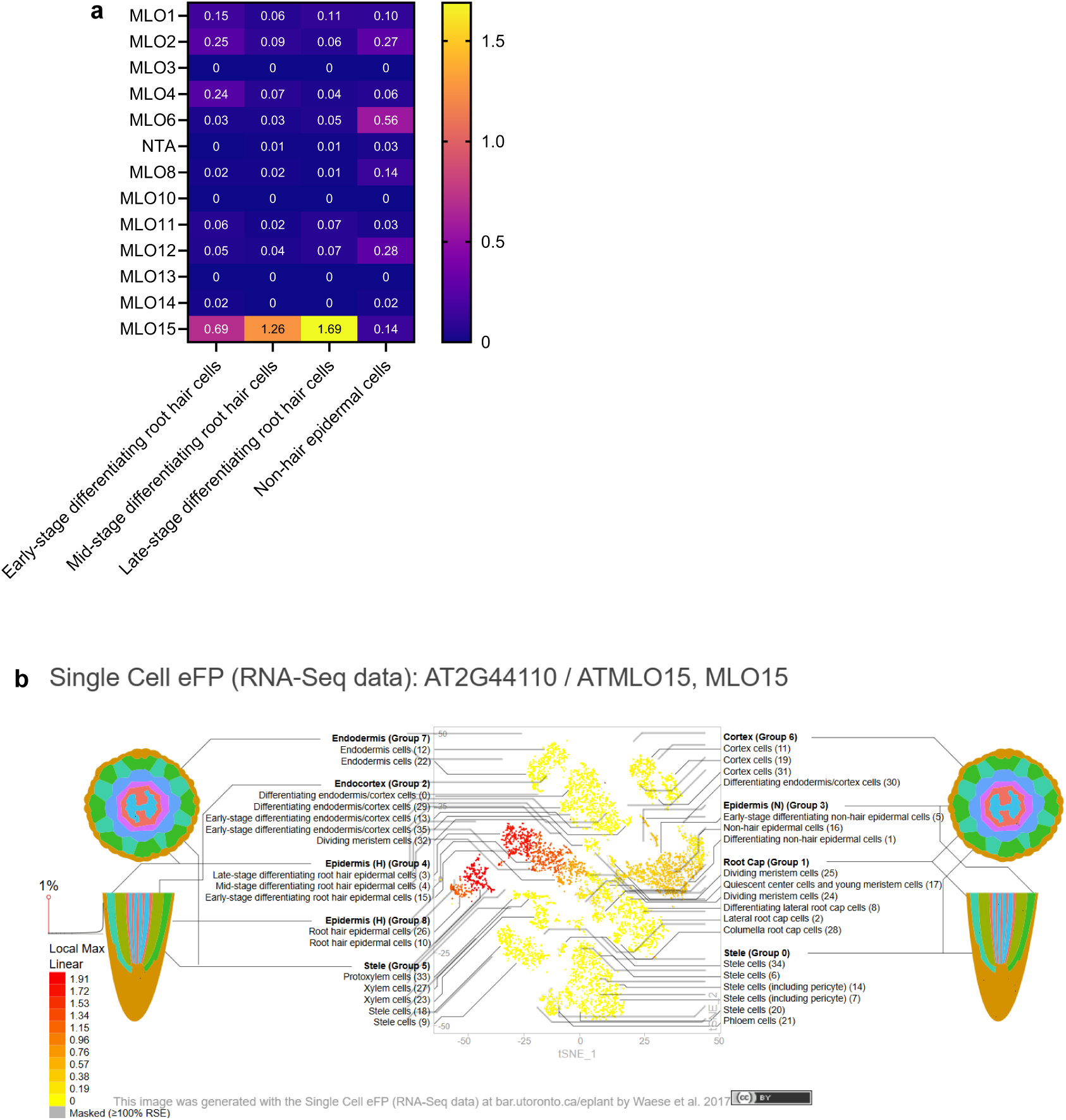
Expression of MLOs in root hair and non-hair epidermal cells. A) Heat map of normalized MLO expression in early, mid, and late-stage differentiating root hair cells and non-hair epidermal cells. Values shown are mean expression from 3 replicates. Expression data for this figure was extracted from the single cell RNA-seq study reported in (Ryu et al., 2019) and visualized as a heat map using GraphPad Prism 10 software. B) MLO15 expression in root cell types. Image was generated with ePlant using single cell RNA-seq data from (Ryu et al., 2019).

**Supplemental Figure 4:**
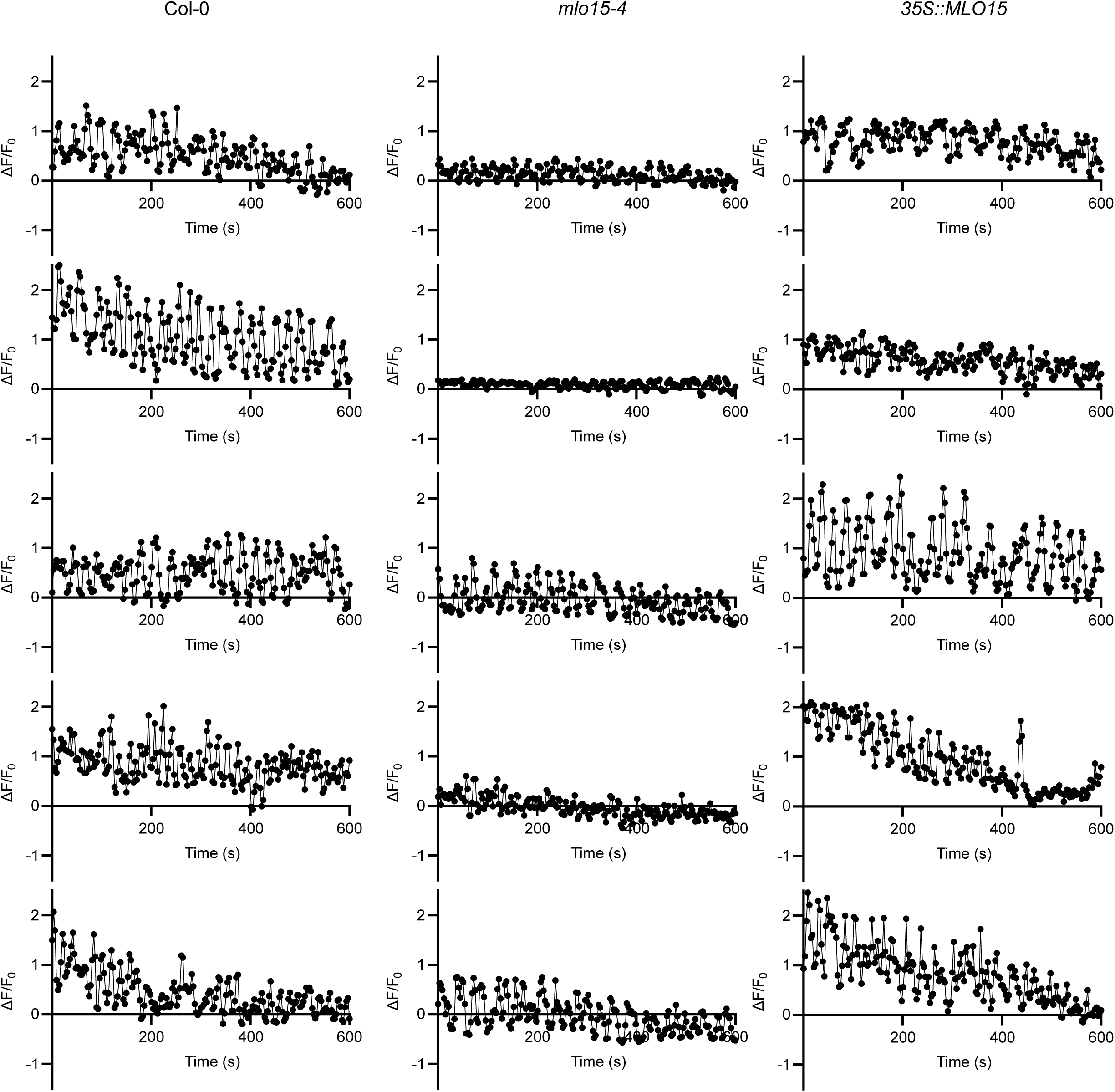
Additional normalized intensity traces of R-GECO1 intensity over 10 minutes in Col-0, *mlo15-4*, and *35S::MLO15*.

**Supplemental Figure 5:**
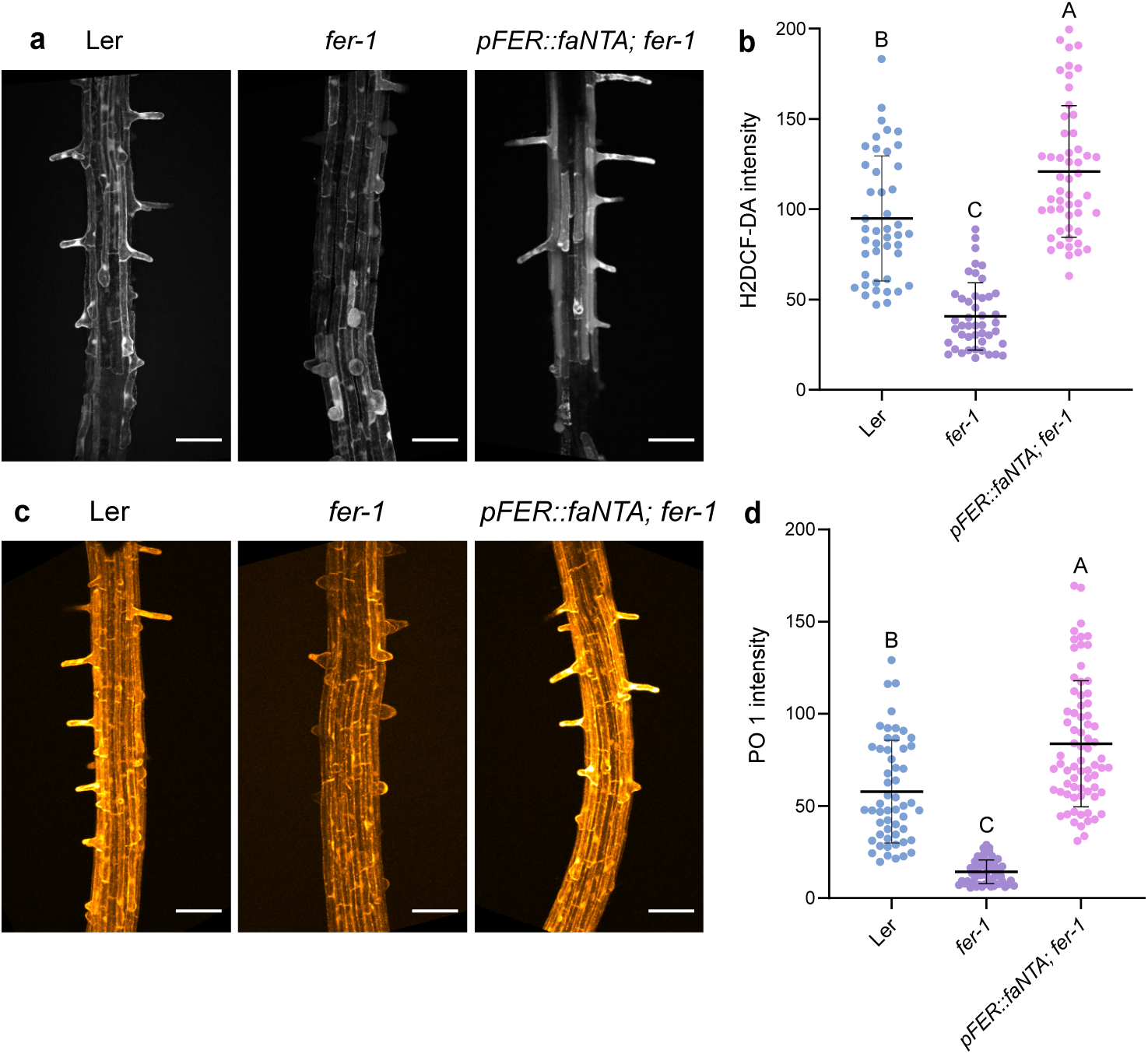
faNTA and positively regulate ROS production in Ler root hairs. A-B) The general ROS indicator H2DCF-DA was used to compare ROS levels in Ler (n=44), *fer-1* (n=45), and *pFER::faNTA;fer-1* (n=52) root hairs. C-D) Peroxy Orange 1 staining was used to quantify H_2_O_2_ levels in Ler (n=54), *fer-1* (n=58), and *pFER::faNTA;fer-1* (n=72) root hairs. Fluorescence intensity was compared in GraphPad Prism 10 with a one-way ANOVA with Tukey test p<0.05. Scale bar = 100 µm.

**Supplemental video 1: R-GECO1 in Col-0 root hairs**

Sum intensity projection of R-GECO1 signal in Col-0 root hairs with images acquired every 3 seconds for 10 minutes. Scale = 100 µm.

**Supplemental video 2: R-GECO1 in *fer-4* root hairs**

Sum intensity projection of R-GECO1 signal in *fer-4* root hairs with images acquired every 3 seconds for 10 minutes. Scale = 100 µm.

**Supplemental video 3: R-GECO1 in *35S::faNTA-YFP;fer-4* root hairs**

Sum intensity projection of R-GECO1 signal in *fer-4* root hairs with images acquired every 3 seconds for 10 minutes. Scale = 100 µm.

**Supplemental video 4: MLO15-GFP expression during root development**

Maximum intensity projection of *pMLO15::MLO15-GFP* imaged every 5 minutes for 4 hours. Scale = 100 µm.

**Supplemental video 5: R-GECO1 in *mlo15-4* root hairs**

Sum intensity projection of R-GECO1 signal in *mlo15-4* root hairs with images acquired every 3 seconds for 10 minutes. Scale = 100 µm.

**Supplemental video 6: R-GECO1 in *35S::MLO15-GFP* root hairs**

Sum intensity projection of R-GECO1 signal in *35S::MLO15-GFP* root hairs with images acquired every 3 seconds for 10 minutes. Scale = 100 µm.

